# Deubiquitinase USP-14 controls intestinal distension-induced immune activation in *Caenorhabditis elegans* via Wnt/β-catenin signaling

**DOI:** 10.1101/2025.11.25.690479

**Authors:** Annesha Ghosh, Jogender Singh

**Affiliations:** Department of Biological Sciences, Indian Institute of Science Education and Research, Mohali, Punjab, 140306, India

**Keywords:** Deubiquitinase, gut distension, innate immunity, USP-14, Wnt signaling

## Abstract

Pathogen infections disrupt multiple host cellular processes, and hosts have consequently evolved mechanisms to detect these perturbations and initiate appropriate immune responses. In *Caenorhabditis elegans*, gut distension caused by bacterial colonization is known to activate innate immunity, yet the molecular mechanisms linking intestinal distension to immune activation remain poorly understood. Here, we perform a forward genetic screen to identify suppressors of intestinal distension-induced immune activation in *C. elegans*. This screen identifies a loss-of-function mutation in the gene encoding the deubiquitinase (DUB) USP-14 as a potent suppressor of immune activation triggered by gut distension. We show that *usp-14* knockout mutants exhibit increased sensitivity to the bacterial pathogens *Pseudomonas aeruginosa* and *Staphylococcus aureus*. Global transcriptomic profiling further demonstrates that USP-14 modulates innate immune responses induced by intestinal distension. Epistasis analyses establish that USP-14 functions through the Wnt/β-catenin signaling pathway to regulate these immune responses. Together, our findings reveal a previously unrecognized role for the conserved DUB USP-14 in host immunity, providing a foundation for future studies investigating how USP-14 modulates immune signaling.

## Introduction

Living in microbe-rich environments, animals have evolved sophisticated strategies to discriminate between harmful and beneficial microorganisms (Blander & Sander, 2012). A central component of this defense is the detection of microbial-associated molecular patterns by conserved pattern-recognition receptors (Akira *et al*, 2006). In parallel, organisms have also evolved mechanisms to sense disruptions in their own cellular physiology, a process termed surveillance immunity (Willmann & Moita, 2025; Pukkila-Worley, 2016). This mode of immune activation is widely conserved across plants, invertebrates, and vertebrates (Cook *et al*, 2015; Pukkila-Worley, 2016; Willmann & Moita, 2025). In *Caenorhabditis elegans*, perturbations of core cellular processes, including translation, endoplasmic reticulum function, mitochondrial function, and the ubiquitin-proteasome system, trigger robust innate immune responses (Melo & Ruvkun, 2012; Pukkila-Worley, 2016; Dunbar *et al*, 2012; McEwan *et al*, 2012; Ghosh & Singh, 2024; Rao *et al*, 2025; Grover *et al*, 2024). Because such perturbations can arise from diverse pathogens, surveillance of cellular integrity likely provides a broad evolutionary advantage (Willmann & Moita, 2025).

In addition to monitoring cell-intrinsic damage, organisms can detect disruptions at the tissue level (Pugin, 2012). For example, epidermal injury in *C. elegans* activates multiple immune pathways, including the p38 MAPK cascade (Taffoni & Pujol, 2015). More recently, intestinal distension caused by pathogenic colonization has been shown to induce innate immune responses in *C. elegans* (Singh & Aballay, 2019a, 2019c; Kumar *et al*, 2019). Remarkably, intestinal distension alone, independent of pathogen exposure, is sufficient to activate immune defenses, indicating that distension is detected as a danger signal (Singh & Aballay, 2019a; Kumar *et al*, 2019). Further studies revealed that this response is mediated by the neuronal acetylcholine receptor ACC-4, which signals to the intestine through the Wnt pathway (Ren *et al*, 2022). Nonetheless, beyond this work, the molecular components that link intestinal distension to immune activation remain poorly defined.

In this study, we conducted a forward genetic screen to identify regulators of immune responses triggered by intestinal distension. Our screen uncovered a loss-of-function mutation in the gene encoding the deubiquitinase (DUB) USP-14, which suppresses distension-induced immune activation and increases susceptibility to *Staphylococcus aureus* and *Pseudomonas aeruginosa*. We showed that intestinal expression of *usp-14* is sufficient to confer immunity to both pathogens. Transcriptomic analyses further revealed that intestinal distension elicits both USP-14-dependent and USP-14-independent innate immune programs. Genetic epistasis experiments demonstrated that USP-14 acts through the canonical Wnt/β-catenin signaling pathway, and notably, USP-14 influences the transcription of multiple Wnt pathway components. Together, our findings identify a critical role for the conserved DUB USP-14 in orchestrating intestinal distension-mediated immune responses through activation of Wnt/β-catenin-dependent innate immunity.

## Results

### The DUB USP-14 is required for immune activation during *C. elegans* intestinal distension

Previous studies have shown that intestinal distension, even in the absence of pathogenic infection, can trigger the activation of innate immune responses (Singh & Aballay, 2019a, 2019c; Kumar *et al*, 2019). Furthermore, it has been reported that a member of the acetylcholine receptor family in the nervous system is required for the activation of immune responses following intestinal distension (Ren *et al*, 2022). To identify additional pathways that mediate the activation of innate immunity upon intestinal distension, we conducted a forward genetic screen using the *aex-5(sa23);clec-60p::gfp* strain. Inhibition of *aex-5*, which encodes a furin-like prohormone convertase, disrupts the defecation motor program, leading to intestinal distension (Branicky & Hekimi, 2006). This distension, in turn, induces the expression of several immune genes, including the C-type lectin gene *clec-60* (Singh & Aballay, 2019a).

The forward genetic screen was designed to isolate mutants exhibiting reduced green fluorescent protein (GFP) expression in *aex-5(sa23);clec-60p::gfp* worms (Figure 1A). We identified four independent mutants with reduced GFP fluorescence (Figure 1B, C). Two of these mutants also showed a marked decrease in the *myo-2p::mCherry* co-injection marker (Figure 1D), suggesting that they might carry mutations causing transgene silencing. The remaining two mutants (#1.5 and #3.2) were backcrossed six times with the parental strain, followed by whole-genome sequencing. Single-nucleotide polymorphism analysis combined with RNA interference (RNAi)-based validation identified *usp-14* and *cec-10* as the causal genes for mutants #1.5 and #3.2, respectively (Figure S1A, B). The *cec-10(jsn20)* allele in mutant #3.2 carried a single G→T substitution mutation, leading to a premature stop codon for glutamic acid (E) codon at 259 position (E259*). Knockdown of *cec-10* led to only a marginal decrease in survival during *P. aeruginosa* infection (Figure S1C). Because *cec-10* lacks a human homolog and has minimal impact on *C. elegans* survival during infection, we did not pursue it further in the context of intestinal distension-induced immune activation.

**Figure 1:**
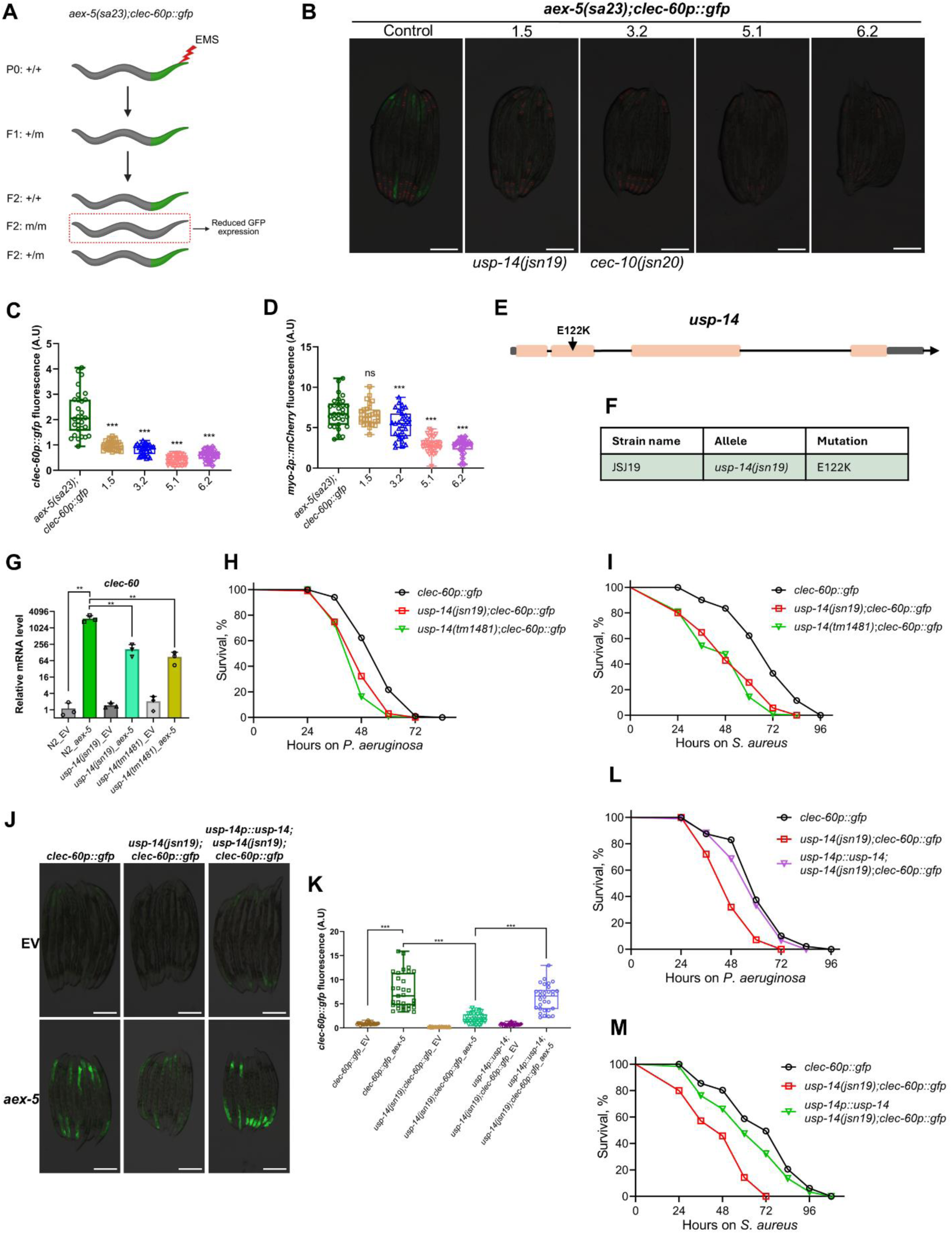
The deubiquitinase USP-14 is required for immune activation during *C. elegans* intestinal distension. (A) Scheme for a forward genetic screen for mutants that have a suppressed *clec-60p::gfp* in *aex-5(sa23);clec-60p::gfp* worms. (B) Representative fluorescence images of the control parental strain *aex-5(sa23);clec-60p::gfp* and the mutants isolated from the forward genetic screen with reduced GFP levels. Scale bar = 200 μm. (C) Quantification of GFP levels of the control parental strain *aex-5(sa23);clec-60p::gfp* and the mutants isolated from the forward genetic screen. \*\*\**p*< 0.001 via ordinary one-way ANOVA followed by Dunnett’s multiple comparisons test (*n* = 30-31 worms each). (D) Quantification of mCherry levels of *myo-2p::mCherry* in the control parental strain *aex-5(sa23);clec-60p::gfp* and the mutants isolated from the forward genetic screen. \*\*\**p* < 0.001 and ns, non-significant via ordinary one-way ANOVA followed by Dunnett’s multiple comparisons test (*n* = 30-31 worms each). (E) Mapping of the *usp-14(jsn19)* allele identified in the forward genetic screen. (F) Summary of the *usp-14(jsn19)* allele. (G) Quantitative reverse transcription-PCR (qRT-PCR) analysis of the immune gene *clec-60* in wild-type N2, *usp-14(jsn19)*, and *usp-14(tm1481)* worms after treatment with the empty vector (EV) control and *aex-5* RNAi. \*\**p*< 0.01 via t-test. Data represents the mean and standard deviation from three independent experiments. (H) Representative survival plots of *clec-60p::gfp*, *usp-14(jsn19);clec-60p::gfp*, and *usp-14(tm1481);clec-60p::gfp* worms on *P. aeruginosa* PA14 at 25°C. \*\*\**p*< 0.001 for the mutants as compared to the control worms (*n* = 105 per condition). (I) Representative survival plots of *clec-60p::gfp*, *usp-14(jsn19);clec-60p::gfp*, and *usp-14(tm1481);clec-60p::gfp* worms on *S. aureus* NCTC832*5* at 25°C. \*\*\**p*< 0.001 for the mutants as compared to the control worms (*n* = 70 for *clec-60p::gfp* and 105 for each *usp-14(jsn19)* and *usp-14(tm1481)*). (J) Representative fluorescence images of *clec-60p::gfp*, *usp-14(jsn19);clec-60p::gfp*, and *usp-14p::usp-14;usp-14(jsn19);clec-60p::gfp* worms exposed to the EV control and *aex-5* RNAi. Scale bar = 200 μm. (K) Quantification of GFP levels of *clec-60p::gfp*, *usp-14(jsn19);clec-60p::gfp*, and *usp-14p::usp-14;usp-14(jsn19);clec-60p::gfp* worms exposed to the EV control and *aex-5* RNAi. \*\*\**p*< 0.001 via t-test (*n* = 30-31 worms each). (L) Representative survival plots of *clec-60p::gfp*, *usp-14(jsn19);clec-60p::gfp*, and *usp-14p::usp-14;usp-14(jsn19);clec-60p::gfp* worms on *P. aeruginosa* PA14 at 25°C. \*\*\**p*< 0.001 between *clec-60p::gfp* and *usp-14(jsn19);clec-60p::gfp*, and non-significant between *clec-60p::gfp* and *usp-14p::usp-14;usp-14(jsn19);clec-60p::gfp* (*n* = 105 per condition). (M) Representative survival plots of *clec-60p::gfp*, *usp-14(jsn19);clec-60p::gfp*, and *usp-14p::usp-14;usp-14(jsn19);clec-60p::gfp* worms on *S. aureus* NCTC832*5* at 25°C. \*\*\**p*< 0.001 between *clec-60p::gfp* and *usp-14(jsn19);clec-60p::gfp*, and \*\**p*< 0.01 between *clec-60p::gfp* and *usp-14p::usp-14;usp-14(jsn19);clec-60p::gfp* (*n* = 105 for *clec-60p::gfp* and *usp-14(jsn19)*, and 60 *for usp-14p::usp-14;usp-14(jsn19);clec-60p::gfp*).

The *usp-14(jsn19)* allele in mutant #1.5 carried a single C→T substitution, resulting in a glutamic acid to lysine (K) change at residue 122 (Figure 1E, F). USP-14 is a cysteine-type DUB that regulates proteasomal protein degradation and is the *C. elegans* homolog of human USP14 (ubiquitin-specific peptidase 14) (Wang *et al*, 2021). To determine whether USP-14 regulates *clec-60* expression at the transcriptional level, we performed quantitative reverse transcription PCR (qRT-PCR) on *usp-14(jsn19)* worms. As expected, RNAi-mediated knockdown of *aex-5* led to robust upregulation of *clec-60* in wild-type N2 worms (Figure 1G). However, *usp-14(jsn19)* worms exhibited significantly attenuated *clec-60* induction, indicating that USP-14 positively regulates *clec-60* transcription. To validate that this effect resulted from *usp-14* loss of function, we examined *clec-60* levels in an independent allele, *usp-14(tm1481)*, which carries a 696 bp deletion encompassing most of the third exon and is likely a null allele. Similar to *usp-14(jsn19)* worms, *usp-14(tm1481)* mutants displayed a significantly reduced induction of *clec-60* upon *aex-5* RNAi compared with wild-type worms (Figure 1G).

We next tested whether USP-14 contributes to host defense against bacterial infection. Both *usp-14(jsn19)* and *usp-14(tm1481)* mutants exhibited reduced survival when exposed to *P. aeruginosa* compared with wild-type worms (Figure 1H). Because *clec-60* expression is also induced by the Gram-positive pathogen *S. aureus* (Irazoqui *et al*, 2010), we examined whether *usp-14* mutants were similarly sensitive to *S. aureus* infection. Consistent with this hypothesis, both *usp-14(jsn19)* and *usp-14(tm1481)* worms showed significantly reduced survival upon *S. aureus* infection (Figure 1I).

To confirm that these phenotypes were specifically due to the loss of USP-14 function, we expressed *usp-14* under its endogenous promoter in *usp-14(jsn19)* mutants. Transgenic expression of *usp-14* restored *clec-60p::gfp* induction upon *aex-5* RNAi (Figure 1J, K) and rescued the survival defects of *usp-14(jsn19)* worms during both *P. aeruginosa* and *S. aureus* infections (Figure 1L, M). Together, these results demonstrate that USP-14 is essential for the activation of innate immune responses triggered by intestinal distension in *C. elegans* and plays a critical role in protecting the host from bacterial infection.

### Intestinal expression of USP-14 is sufficient to activate immunity during intestinal distension

We next examined the spatial expression pattern of *usp-14* in *C. elegans*. To this end, we generated transgenic worms expressing GFP under the control of the *usp-14* promoter (*usp-14p::gfp*). GFP fluorescence was primarily detected in a subset of neuronal cells (Figure 2A). Based on this expression pattern, we hypothesized that neuronal *usp-14* might be required for regulating innate immune responses. To test this, we expressed *usp-14* under the pan-neuronal promoter *rgef-1p* in *usp-14(jsn19)* mutants and assessed both *clec-60* expression and survival upon bacterial infection. Surprisingly, neuronal expression of *usp-14* failed to restore *clec-60* expression (Figure 2B, C) or improve the survival of *usp-14(jsn19)* worms upon bacterial infection (Figure 2D, E).

**Figure 2:**
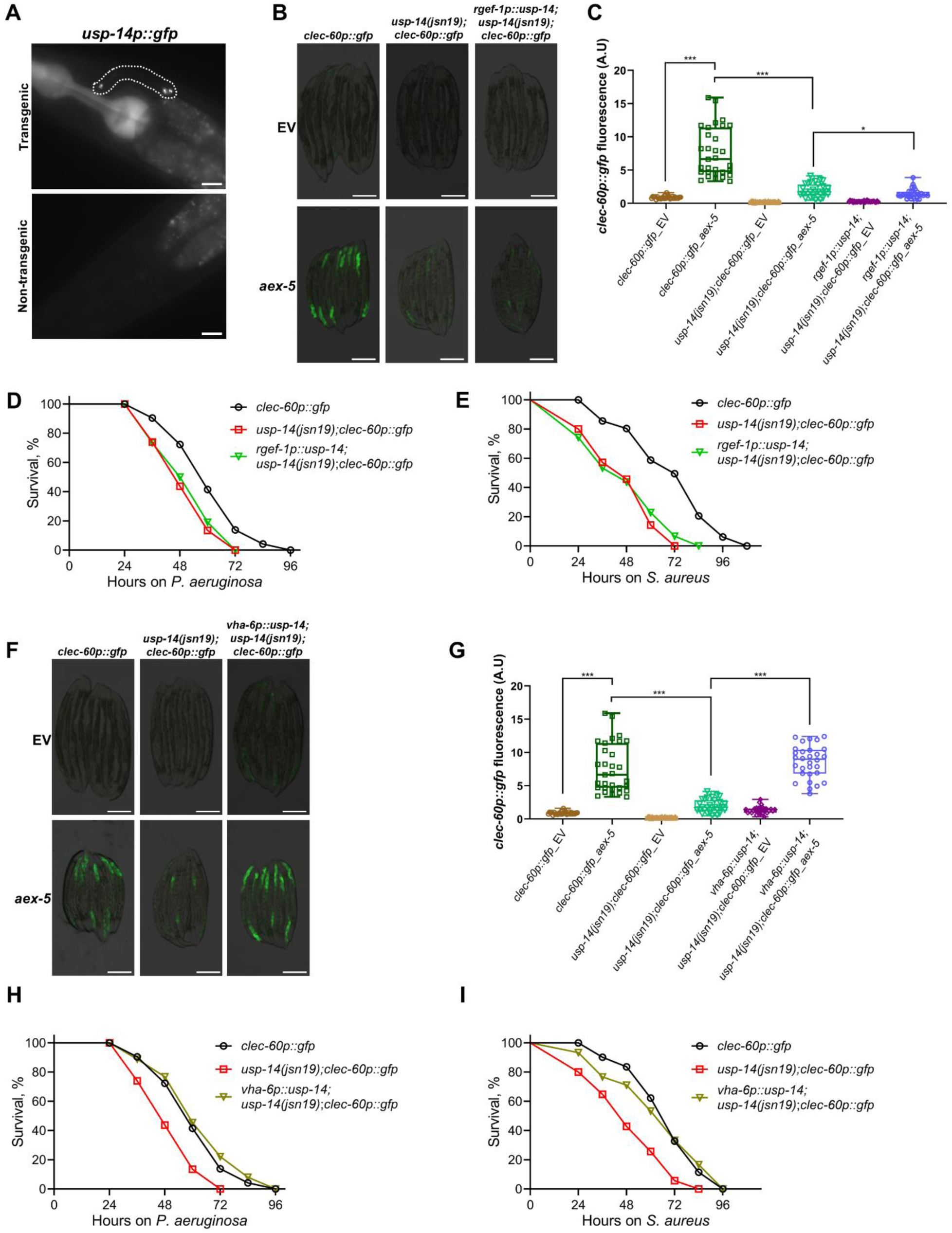
Intestinal expression of USP-14 is sufficient to activate immunity during intestinal distension. (A) Representative fluorescence images of *usp-14p::gfp* worm showing GFP in some head neurons (highlighted by the dotted line) along with a control non-transgenic worm. The fluorescence in the pharynx in the transgenic worm is from *myo-2p::mCherry*. Scale bar = 10 μm. (B) Representative fluorescence images of *clec-60p::gfp*, *usp-14(jsn19);clec-60p::gfp*, and *rgef-1p::usp-14;usp-14(jsn19);clec-60p::gfp* worms exposed to the empty vector (EV) control and *aex-5* RNAi. Scale bar = 200 μm. (C) Quantification of GFP levels of *clec-60p::gfp*, *usp-14(jsn19);clec-60p::gfp*, and *rgef-1p::usp-14;usp-14(jsn19);clec-60p::gfp* worms exposed to the EV control and *aex-5* RNAi. \*\*\**p*< 0.001 and \**p*< 0.05 via unpaired t-test (*n* = 30-31 worms each). The controls *clec-60p::gfp* and *usp-14(jsn19);clec-60p::gfp* for this experiment are the same as shown in Figures 1K and 2G. (D) Representative survival plots of *clec-60p::gfp*, *usp-14(jsn19);clec-60p::gfp*, and *rgef-1p::usp-14;usp-14(jsn19);clec-60p::gfp* worms on *P. aeruginosa* PA14 at 25°C. \*\*\**p*< 0.001 between *clec-60p::gfp* and *usp-14(jsn19);clec-60p::gfp*, and non-significant between *usp-14(jsn19);clec-60p::gfp* and *rgef-1p::usp-14;usp-14(jsn19);clec-60p::gfp* (*n* = 105 per condition). (E) Representative survival plots of *clec-60p::gfp*, *usp-14(jsn19);clec-60p::gfp*, and *rgef-1p::usp-14;usp-14(jsn19);clec-60p::gfp* worms on *S. aureus* NCTC832*5* at 25°C. \*\*\**p*< 0.001 between *clec-60p::gfp* and *usp-14(jsn19);clec-60p::gfp*, and non-significant between *usp-14(jsn19);clec-60p::gfp* and *rgef-1p::usp-14;usp-14(jsn19);clec-60p::gfp* (*n* = 105 per condition). The curves shown for *clec-60p::gfp* and *usp-14(jsn19);clec-60p::gfp* on *S. aureus* NCTC8325 are the same as shown in Figure 1M. (F) Representative fluorescence images of *clec-60p::gfp*, *usp-14(jsn19);clec-60p::gfp*, and *vha- 6p::usp-14;usp-14(jsn19);clec-60p::gfp* worms exposed to the EV control and *aex-5* RNAi. Scale bar = 200 μm. (G) Quantification of GFP levels of *clec-60p::gfp*, *usp-14(jsn19);clec-60p::gfp*, and *vha-6p::usp-14;usp-14(jsn19);clec-60p::gfp* worms exposed to the EV control and *aex-5* RNAi. \*\*\**p*< 0.001 via t-test (*n* = 30-31 worms each). The controls *clec-60p::gfp* and *usp-14(jsn19);clec-60p::gfp* for this experiment are the same as shown in Figures 1K and 2C. (H) Representative survival plots of *clec-60p::gfp*, *usp-14(jsn19);clec-60p::gfp*, and *vha-6p::usp-14;usp-14(jsn19);clec-60p::gfp* worms on *P. aeruginosa* PA14 at 25°C. \*\*\**p*< 0.001 between *clec-60p::gfp* and *usp-14(jsn19);clec-60p::gfp*, and non-significant between *clec-60p::gfp* and *vha-6p::usp-14;usp-14(jsn19);clec-60p::gfp* (*n* = 105 per condition). The curves shown for *clec-60p::gfp* and *usp-14(jsn19);clec-60p::gfp* on *P. aeruginosa* PA14 are the same as shown in Figure 2D. (I) Representative survival plots of *clec-60p::gfp*, *usp-14(jsn19);clec-60p::gfp*, and *vha-6p::usp-14;usp-14(jsn19);clec-60p::gfp* worms on *S. aureus* NCTC832*5* at 25°C. \*\*\**p*< 0.001 between *clec-60p::gfp* and *usp-14(jsn19);clec-60p::gfp*, and non-significant between *clec-60p::gfp* and *vha-6p::usp-14;usp-14(jsn19);clec-60p::gfp* (*n* = 70 for *clec-60p::gfp*, 105 for *usp-14(jsn19);clec-60p::gfp*, and 90 for *vha-6p::usp-14;usp-14(jsn19);clec-60p::gfp).* The curves shown for *clec-60p::gfp* and *usp-14(jsn19);clec-60p::gfp* on *S. aureus* NCTC8325 are the same as shown in Figure 1I.

Given that *clec-60* expression is primarily localized to intestinal cells, the main immune barrier against bacterial pathogens in *C. elegans*, we next asked whether USP-14 functions in the intestine to mediate immune activation. To address this, we expressed *usp-14* under the intestine-specific promoter *vha-6p* in *usp-14(jsn19)* mutants and evaluated both *clec-60* induction and pathogen resistance. Remarkably, intestinal expression of *usp-14* completely rescued *clec-60* induction upon *aex-5* RNAi (Figure 2F, G). Consistently, intestinal expression also restored the survival of *usp-14(jsn19)* worms during both *P. aeruginosa* and *S. aureus* infections (Figure 2H, I). Together, these findings demonstrate that intestinal, but not neuronal, expression of USP-14 is sufficient to activate innate immune responses during intestinal distension in *C. elegans*.

### Intestinal distension induces USP-14-dependent and -independent innate immune response genes

To elucidate the role of USP-14 in intestinal distension-mediated immune activation, we performed transcriptomic profiling by RNA sequencing. We compared the global gene expression patterns of wild-type N2 and *usp-14(tm1481)* worms grown to adulthood on the empty vector (EV) control or *aex-5* RNAi (Table S1 and S2).

We first examined whether USP-14 influences the transcriptome under basal conditions, in the absence of intestinal distension. Comparison of *usp-14(tm1481)* mutants and wild-type worms grown on EV revealed 127 differentially expressed genes, of which 86 were upregulated and 41 were downregulated (Figure 3A, Table S3). Gene Ontology (GO) enrichment analysis of both upregulated (Figure 3B) and downregulated (Figure 3C) genes indicated a strong overrepresentation of innate immune response genes. These results suggested that loss of *usp-14* perturbs immune homeostasis, potentially leading to constitutive activation of MAP kinase signaling pathways (Figure S2A, B).

**Figure 3:**
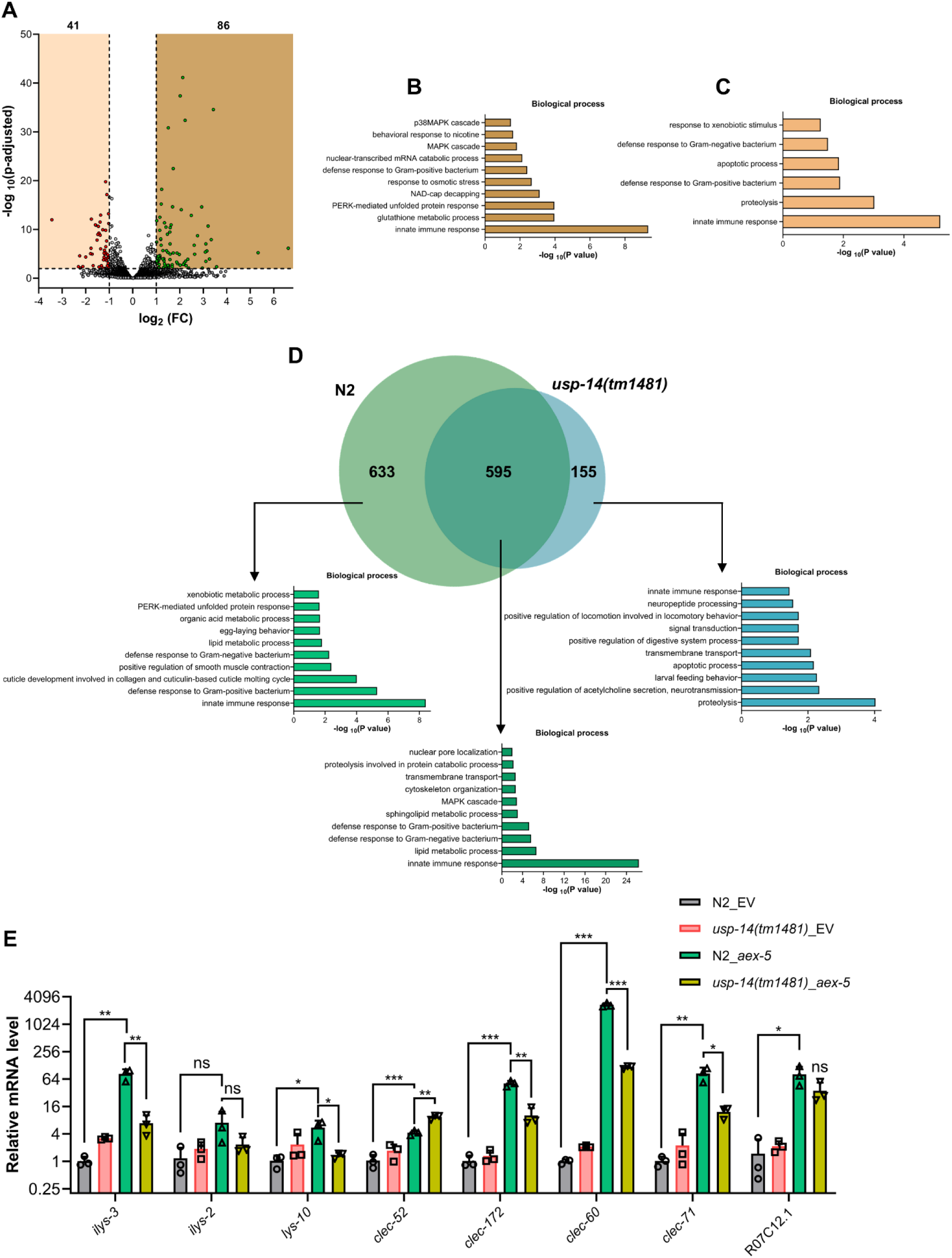
Intestinal distension induces USP-14-dependent and -independent innate immune response genes. (A) Volcano plot of upregulated and downregulated genes in *usp-14(tm1481)* worms compared to wild-type N2 worms grown on empty vector (EV) control RNAi. Green and red dots represent significantly upregulated and downregulated genes, respectively, while the gray dots represent the genes that are not differentially regulated. (B)-(C) Gene Ontology (GO) enrichment analysis for upregulated (B) and downregulated (C) genes in *usp-14(tm1481)* worms compared to wild-type N2 worms grown on the EV control RNAi. (D) Venn diagram showing the overlap between genes upregulated upon *aex-5* RNAi in N2 and *usp-14(tm1481)* worms. The GO analysis for the biological processes of unique and common genes is shown. (E) qRT-PCR analysis of innate immune response genes in wild-type N2 and *usp-14(tm1481)* worms after treatment with the EV control and *aex-5* RNAi. \*\*\**p*< 0.001, \*\**p*< 0.01, \**p*< 0.05, ns, non-significant via t-test. Data represent the mean and standard deviation from three independent experiments.

Intestinal distension is known to activate innate immune responses (Singh & Aballay, 2019a; Kumar *et al*, 2019). Therefore, we analyzed whether *aex-5* RNAi in N2 worms also led to the induction of innate immune response genes. The knockdown of *aex-5* resulted in the upregulation of 1241 genes and downregulation of 209 genes (Table S1). As expected, the upregulated genes were strongly enriched in innate immune responses (Figure S2C). We next investigated which of these *aex-5* RNAi-induced genes required USP-14 for their activation. A comparative analysis between *aex-5* RNAi-treated N2 and *usp-14(tm1481)* worms identified 633 genes whose induction was USP-14-dependent (Figure 3D, Table S4). GO enrichment analysis revealed that these USP-14-dependent genes were highly enriched for innate immune response functions (Figure 3D).

Because USP-14 also regulated a subset of immune genes under basal conditions (Figure 3C), we examined whether the USP-14-dependent genes induced during intestinal distension overlap with those regulated in the absence of distension. Notably, no overlap was observed between the genes downregulated in *usp-14(tm1481)* mutants under basal conditions and those dependent on USP-14 during intestinal distension (Figure S2D). This finding indicated that USP-14 specifically regulates a distinct set of immunity genes upon intestinal distension, beyond its basal role in immune homeostasis.

To validate the RNA sequencing results, we performed qRT-PCR on a subset of immunity genes that were upregulated in N2 worms but not in *usp-14(tm1481)* mutants following *aex-5* RNAi treatment (Figure 3E). Most of these genes showed significantly reduced expression in *usp-14(tm1481)* worms compared to N2 worms under *aex-5* RNAi, confirming the transcriptomic results. We then analyzed the GO enrichment for the 595 genes that were upregulated in both N2 and *usp-14(tm1481)* worms upon *aex-5* RNAi. These genes were also strongly enriched in innate immune response terms (Figure 3D), whereas genes uniquely upregulated in *usp-14(tm1481)* mutants showed no such enrichment. Collectively, these findings demonstrate that intestinal distension triggers both USP-14-dependent and USP-14-independent innate immune response pathways in *C. elegans*, highlighting a dual regulatory mechanism underlying distension-induced immune activation.

### USP-14 modulates immunity via the Wnt/β-catenin signaling pathway

We next investigated whether USP-14 regulates intestinal distension-mediated immune activation through known immune signaling pathways. To this end, we generated several double mutants in the *clec-60p::gfp* background, including *hlh-30(tm1978);usp-14(tm1481)*, *nhr-49(nr2041);usp-14(tm1481)*, and *pmk-1(km25);usp-14(tm1481)*. The TFEB ortholog HLH-30, the nuclear hormone receptor NHR-49, and the NSY-1/SEK-1/PMK-1 MAP kinase cascade are all known to mediate host defense against *P. aeruginosa* and *S. aureus* infections (Kim *et al*, 2002; Naim *et al*, 2021; Rao & Singh, 2025; Sifri *et al*, 2003; Wani *et al*, 2021), similar to USP-14 as shown in this study.

RNAi-mediated knockdown of *aex-5* resulted in robust induction of *clec-60p::gfp* in *hlh-30(tm1978)*, *nhr-49(nr2041)*, and *pmk-1(km25)* mutants (Figure S3A, B). Likewise, exposure of these mutants to *S. aureus* infection also induced *clec-60p::gfp* expression (Figure S4A, B). However, this induction was strongly suppressed when these mutations were combined with *usp-14(tm1481)*, both under *aex-5* RNAi and during *S. aureus* infection (Figures S3, S4). These findings indicated that USP-14 regulates intestinal distension-mediated activation of *clec-60* independently of HLH-30, NHR-49, and PMK-1. Furthermore, double mutants of *usp-14(tm1481)* with *hlh-30(tm1978)*, *nhr-49(nr2041)*, or *pmk-1(km25)* exhibited enhanced susceptibility to *P. aeruginosa* and *S. aureus* infections compared to either single mutant (Figure S5A-F), further supporting that USP-14 acts independently of these canonical immune pathways.

We next investigated whether USP-14-mediated immune regulation is associated with the Wnt/β-catenin signaling pathway. The β-catenin homolog BAR-1 is known to regulate innate immunity in *C. elegans* intestinal cells (Irazoqui *et al*, 2008; Labed *et al*, 2018), and intestinal distension has been shown to activate immune responses through components of the Wnt pathway (Ren *et al*, 2022). To test whether USP-14 acts through this pathway, we first confirmed that knockdown of *bar-1* suppressed intestinal distension-induced *clec-60* expression (Figure 4A, B). Knockdown of *bar-1* did not further reduce expression of *clec-60* in the *usp-14(jsn19)* background. Notably, while *bar-1* knockdown increased the susceptibility of N2 worms to *P. aeruginosa* and *S. aureus*, it did not further enhance the sensitivity of *usp-14(tm1481)* worms (Figure 4C-F), suggesting that USP-14 and BAR-1 function in the same pathway.

**Figure 4:**
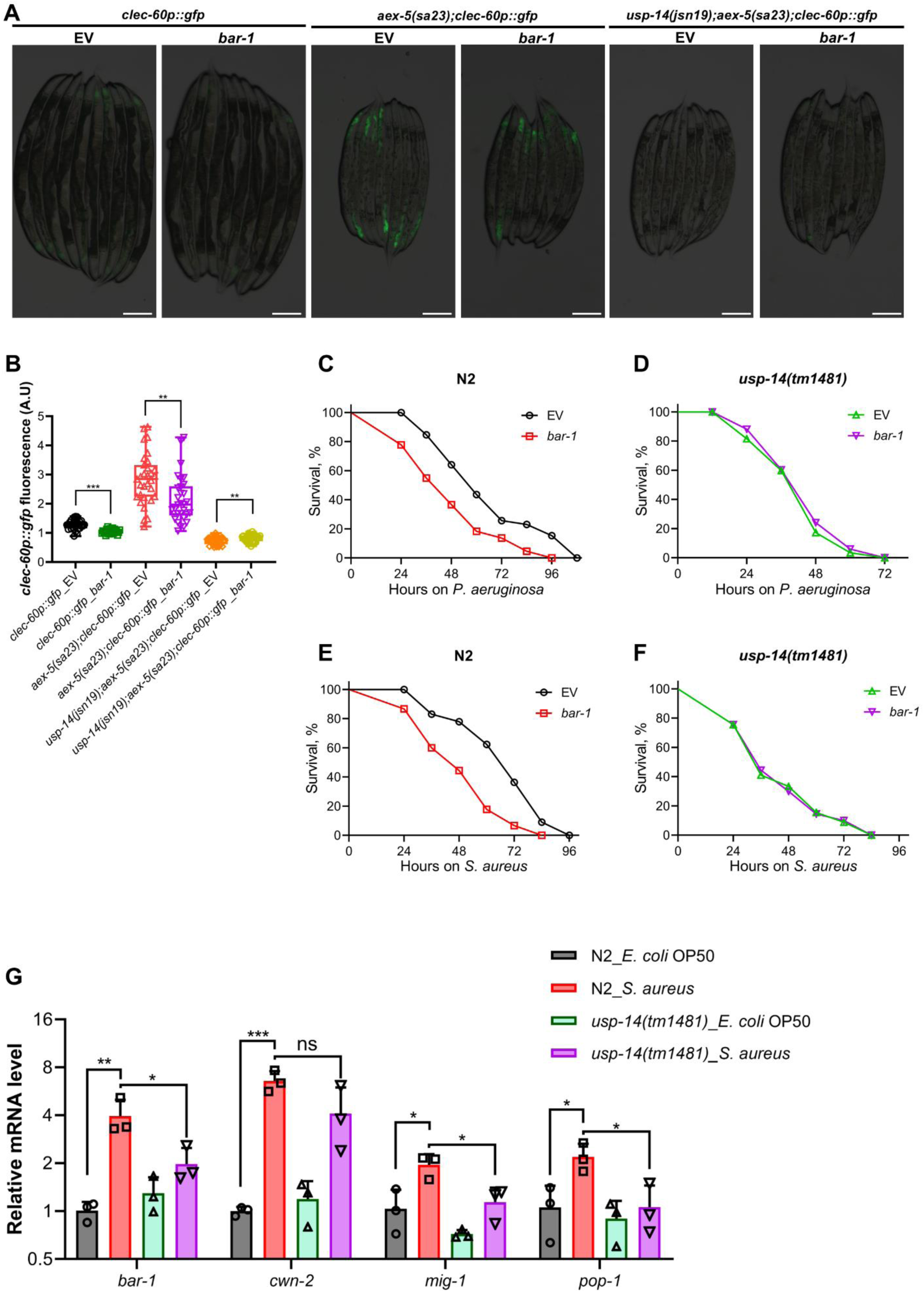
USP-14 modulates immunity via the Wnt/β-catenin signaling pathway. (A) Representative fluorescence images of *clec-60p::gfp*, *aex-5(sa23);clec-60p::gfp*, and *usp-14(jsn19);aex-5(sa23);clec-60p::gfp* worms exposed to the empty vector (EV) control and *bar-1* RNAi. Scale bar = 200 μm. (B) Quantification of GFP levels of *clec-60p::gfp*, *aex-5(sa23);clec-60p::gfp*, and *usp-14(jsn19);aex-5(sa23);clec-60p::gfp* worms exposed to the EV control and *bar-1* RNAi. \*\*\**p*< 0.001 and \*\**p*< 0.01 via unpaired t-test (*n* = 28-31 worms each). (C) Representative survival plots of control N2 worms exposed to the EV and *bar-1* RNAi on *P. aeruginosa* PA14 at 25°C. \*\*\**p*< 0.001 (*n* = 60 on EV and 90 on *bar-1* RNAi). (D) Representative survival plots of *usp-14(tm1481)* worms exposed to the EV and *bar-1* RNAi on *P. aeruginosa* PA14 at 25°C. ns, non-significant (*n* = 105 per condition). (E) Representative survival plots of control N2 worms exposed to the EV and *bar-1* RNAi on *S. aureus* NCTC8325 at 25°C. \*\*\**p*< 0.001 (*n* = 90 per condition). (F) Representative survival plots of control *usp-14(tm1481)* worms exposed to the EV and *bar-1* RNAi on *S. aureus* NCTC8325 at 25°C. ns, non-significant (*n* = 90 per condition). (G) qRT-PCR analysis of the Wnt/β-catenin signaling pathway genes in wild-type N2 and *usp-14(tm1481)* worms after 12 hours of infection with *S. aureus* NCTC8325, compared with *E. coli* OP50 controls at 25 °C. \*\*\**p*< 0.001, \*\**p*< 0.01, \**p*< 0.05, and ns, non-significant via t-test. Data represent the mean and standard deviation from three independent experiments.

Previous studies have shown that *S. aureus* infection induces the expression of *bar-1* (encoding the transcriptional effector of Wnt signaling), *pop-1* (encoding the TCF transcription factor), and other Wnt pathway components such as *mig-1* (encoding a Wnt receptor) and *cwn-2* (encoding a Wnt ligand) (Labed *et al*, 2018; Ren *et al*, 2022). To determine whether USP-14 regulates Wnt signaling at the transcriptional level, we quantified the mRNA levels of these genes upon *S. aureus* infection. The results revealed that *S. aureus*-induced expression of Wnt signaling pathway genes was significantly reduced in *usp-14(tm1481)* worms compared to wild type (Figure 4G). Together, these findings demonstrate that USP-14 modulates innate immune responses by regulating the transcriptional activation of multiple components of the Wnt/β-catenin signaling pathway, positioning USP-14 as an upstream regulator of Wnt-mediated immunity during intestinal distension in *C. elegans*.

## Discussions

Recent work across multiple organisms has demonstrated that intestinal distension profoundly influences food-seeking behavior, immunity, and lifespan (Das *et al*, 2024; Min *et al*, 2021; Duvall *et al*, 2019; Kumar *et al*, 2019; Singh & Aballay, 2019b; Wang *et al*, 2025). In *C. elegans*, gut distension modulates pathogen avoidance by activating defined neuroendocrine pathways (Filipowicz *et al*, 2021; Singh & Aballay, 2019a, 2019c). These distension-induced avoidance behaviors can also be inherited by progeny through histone H4 Lys8 acetylation in the germline (Hong *et al*, 2021). Beyond behavioral outputs, intestinal distension also initiates robust innate immune activation (Singh & Aballay, 2019a; Kumar *et al*, 2019). This response is mediated by the neuronal acetylcholine receptor ACC-4, which signals to the intestine via the Wnt/β-catenin pathway (Ren *et al*, 2022). In this study, we identify the DUB USP-14 as an intestinal factor required for this distension-mediated immune activation. Both ACC-4 and USP-14 converge on the Wnt/β-catenin axis; however, it remains unclear whether they function within a single pathway or act in parallel to control Wnt-dependent immune gene expression. Moreover, because intestinal distension also activates USP-14-independent immune responses, additional signaling mechanisms must operate downstream of gut distension.

USP14 is a highly conserved proteasome-associated DUB with dual activities in regulating protein stability. It can inhibit proteasomal degradation by trimming ubiquitin chains from substrates bound to the proteasome (Lee *et al*, 2016, 2010), yet it can also promote protein degradation by facilitating 20S proteasome gate opening (Peth *et al*, 2009). In mammals, USP14 has been implicated in cancer progression by stabilizing oncogenic factors such as hypoxia-inducible factor and Jun kinase (Du *et al*, 2023; Lv *et al*, 2021), and it has become a therapeutic target in several malignancies (Tian *et al*, 2014; Morgan *et al*, 2023). USP14 can also suppress T-cell function, thereby contributing to immune evasion in tumors (Shi *et al*, 2022; Yuan *et al*, 2025). Beyond cancer, USP14 also negatively regulates RIG-I-dependent antiviral immunity (Li *et al*, 2019). Conversely, USP14 activity in lung epithelial cells promotes cytokine release by enhancing I-κB degradation (Mialki *et al*, 2013). Our findings add to this complex landscape by demonstrating that USP-14 is required for the activation of innate immune responses triggered by intestinal distension in *C. elegans*. Thus, the immunomodulatory functions of USP14 appear to be strongly context dependent.

USP14 has also been linked to the Wnt/β-catenin pathway at multiple mechanistic levels. In mammalian cells, USP14 enhances Wnt signaling by deubiquitinating Dishevelled (Jung *et al*, 2013). Ubiquitin immunoprecipitation studies in USP14 inactive mutants have further identified β-catenin as a USP14 substrate (Chadchankar *et al*, 2019), and subsequent work demonstrated that USP14-mediated stabilization of β-catenin promotes diffuse large B-cell lymphoma proliferation (Wang *et al*, 2024). Consistently, curcumin-mediated inhibition of cancer progression has been attributed, in part, to modulation of β-catenin through USP-14 (Shen *et al*, 2025). These findings collectively highlight the evolutionarily conserved connection between USP14 and Wnt/β-catenin signaling.

Our data now extend this relationship to innate immunity in *C. elegans*, showing that USP-14 regulates the transcriptional activation of Wnt/β-catenin pathway components during intestinal distension. Interestingly, the mode of USP-14 action in *C. elegans* appears distinct from previously described post-translational regulation of Wnt components in mammalian systems. The ability of USP-14 to influence Wnt signaling at the transcriptional level suggests the existence of additional substrates or interacting proteins that modulate transcriptional outputs downstream of intestinal distension. In future work, it will be important to identify the direct USP-14 substrates that enable transcriptional activation of Wnt/β-catenin signaling and to determine how these molecular mechanisms interface with neuronal ACC-4 signaling and other distension-responsive pathways.

## Materials and methods

### Bacterial strains

The following bacterial strains were used in this study: *Escherichia coli* OP50, *E. coli* HT115(DE3), *Pseudomonas aeruginosa* PA14, and *Staphylococcus aureus* NCTC8325. Cultures of *E. coli* OP50, *E. coli* HT115(DE3), and *P. aeruginosa* PA14 were grown in Luria-Bertani (LB) broth at 37°C, while *S. aureus* NCTC8325 was cultured in tryptic soy (TS) broth at 37°C.

### *C. elegans* strains and growth conditions

*C. elegans* hermaphrodites were maintained at 20°C on nematode growth medium (NGM) plates seeded with *E. coli* OP50 as the food source, unless stated otherwise. The following strains were used in the study: JIN810 *clec-60p::gfp*, JT23 *aex-5(sa23)*, *usp-14(tm1481)*, KU25 *pmk-1(km25)*, JIN1375 *hlh-30(tm1978),* STE68 *nhr-49(nr2041).* The *aex-5(sa23);clec-60p::gfp*, *usp-14(jsn19);clec-60p::gfp, usp-14(tm1481);clec-60p::gfp*, *pmk-1(km25);clec-60p::gfp, pmk-1(km25);usp-14(tm1481);clec-60p::gfp, hlh-30(tm1978);clec-60p::gfp, hlh-30(tm1978);usp-14(tm1481);clec-60p::gfp, nhr-49(nr2041);clec-60p::gfp*, and *nhr-49(nr2041);usp-14(tm1481);clec-60p::gfp* strains were obtained by standard genetic crosses. The following strains were generated in this study: JSJ19 *usp-14(jsn19),* JSJ20 *cec-10(jsn20)*, *usp-14(jsn19);clec-60p::gfp;jsnEx6[usp-14p::usp-14 + myo-3p::mCherry], usp-14(jsn19);clec-60p::gfp;jsnEx7[rgef-1p::usp-14 + myo-3p::mCherry], usp-14(jsn19);clec-60p::gfp;jsnEx8[vha-6p::usp-14 + myo-3p::mCherry],* and *jsnEx9[usp-14p::gfp + myo-2p::mCherry].* Some of the strains were obtained from the Caenorhabditis Genetics Center (University of Minnesota, Minneapolis, MN). The *usp-14(tm1481)* allele was obtained from the National BioResource Project (NBRP), Japan.

### RNA interference (RNAi)

RNAi was used to induce gene-specific loss-of-function phenotypes by feeding *C. elegans* with *E. coli* HT115(DE3) strains expressing double-stranded RNA corresponding to the target gene. RNAi experiments were performed as described previously (Ghosh & Singh, 2024). Briefly, *E. coli* HT115(DE3) carrying the appropriate RNAi plasmids were grown overnight in LB broth containing ampicillin (100 µg/mL) at 37°C overnight on a shaker, concentrated 10 times, and plated onto RNAi NGM plates containing 100 µg/mL ampicillin and 3 mM isopropyl β-D-thiogalactoside. The plated bacteria were allowed to grow overnight at 37°C. Gravid adults were transferred to RNAi-expressing bacterial lawns and allowed to lay eggs for 2 hours. Adults were then removed, and the eggs were incubated at 20°C for 72 hours before being used for downstream assays. All RNAi clones were obtained from the Ahringer RNAi library and verified by sequencing. For *bar-1* RNAi experiments, worms were grown on empty vector-containing *E. coli* until the L4 stage and subsequently transferred to *bar-1* RNAi plates.

### Forward genetic screen for mutants with suppressed *clec-60p::gfp* expression

Ethyl methanesulfonate (EMS) mutagenesis screens were performed using the *aex-5(sa23);clec-60p::gfp* strain as described previously (Singh, 2021). Approximately 2,500 synchronized late L4 larvae were treated with 50 mM EMS for 4 hours, followed by three washes with M9 buffer. The EMS-treated animals (P0 generation) were transferred to 9-cm NGM plates seeded with *E. coli* OP50 and allowed to lay eggs (F1 progeny) overnight. Subsequently, the P0 animals were washed away with M9 medium, while the F1 eggs remained attached to the bacterial lawn. The F1 progeny were allowed to develop into adults, which were then bleached to obtain F2 eggs. F2 eggs were transferred to *E. coli* OP50-seeded NGM plates and incubated at 20°C for 96 hours. Plates were screened for worms exhibiting suppressed *clec-60p::gfp* expression levels. A total of ∼50,000 haploid genomes were screened, resulting in the isolation of four mutants with reduced GFP expression. Each mutant was backcrossed six times with the parental *aex-5(sa23);clec-60p::gfp* strain before further analysis.

### Whole-genome sequencing (WGS) and data analysis

Genomic DNA was isolated following previously described methods (Gokul & Singh, 2022; Ravi *et al*, 2023). Mutant worms were cultivated at 20°C on NGM plates seeded with *E. coli* OP50 until starvation. For each strain, animals from four 9-cm plates were collected using M9 buffer, washed three times, and incubated in M9 with gentle rotation for 2 hours to clear intestinal contents. The worms were washed three additional times with distilled water and stored at -80°C until DNA extraction. Genomic DNA was extracted using the Gentra Puregene Kit (Qiagen, Netherlands). DNA libraries were prepared according to the standard Illumina protocol (San Diego, CA). Whole-genome sequencing was performed on an Illumina NovaSeq 6000 platform utilizing 150 paired-end nucleotide reads. Library preparation and sequencing were carried out at the National Genomics Core, National Institute of Biomedical Genomics, Kalyani, India.

Whole-genome sequence data were analyzed using the Galaxy web platform, as described earlier (Gokul & Singh, 2022). Briefly, the forward and reverse FASTQ reads, *C. elegans* reference genome Fasta file (ce11M.fa), and the gene annotation file (SnpEff4.3 WBcel235.86) were input into the Galaxy workflow. Low-quality bases were trimmed using the Sickle tool, and the filtered reads were aligned to the reference genome using BWA-MEM. Duplicate reads mapping to multiple genomic locations were removed using MarkDuplicates. Variant calling was performed using FreeBayes, which identifies small polymorphisms, including single-nucleotide polymorphisms (SNPs), insertions and deletions (indels), multi-nucleotide polymorphisms (MNPs), and complex variants smaller than the typical short-read alignment length. Shared variants among different mutants were filtered out to identify unique mutations. Genetic variants were annotated using the SnpEff 4.3 WBcel235.86 database to predict their effects, including amino acid substitutions. Finally, linkage maps for each mutant were generated based on the identified variant profiles.

### C. elegans killing assays on P. aeruginosa PA14

Full lawn killing assays of *C. elegans* on *P. aeruginosa* PA14 were performed as previously described (Ghosh & Singh, 2024; Das *et al*, 2024). Briefly, single colonies of *P. aeruginosa* PA14 were inoculated into 2 mL of LB broth and incubated at 37°C with shaking for 10-12 hours. Bacterial lawns were prepared by spreading 20 µL of the culture onto modified NGM agar plates (3.5 cm diameter) containing 0.35% peptone. The seeded plates were incubated at 37°C for 10-12 hours and then cooled to room temperature for at least 30 minutes before use. After that, synchronized 1-day-old adult worms were transferred to the *P. aeruginosa*-seeded plates and maintained at 25°C. Live animals were transferred daily to freshly prepared *P. aeruginosa* plates to avoid confounding effects of progeny. Worms were examined at the indicated time points and scored as dead if they failed to respond to gentle prodding. For killing assays involving *bar-1* RNAi, worms were grown on empty vector-containing *E. coli* until the L4 stage and subsequently transferred to *bar-1* RNAi plates. The 1-day-old adult worms were then transferred to *P. aeruginosa*-seeded plates and maintained at 25°C. Each experimental condition was assayed in at least three independent biological replicates. Detailed statistical analyses for all survival assays are provided in Table S5.

### C. elegans killing assays on S. aureus

Individual *S. aureus* colonies were inoculated into 2 mL of TS broth and cultured overnight at 37°C with shaking. The overnight culture was diluted 1:10 in fresh TS broth, and 20 µL of the diluted suspension was spread onto 3.5 cm NGM plates. The plates were incubated at 37°C for 6 hours to allow lawn formation, then cooled to room temperature for at least 30 minutes before use. After that, synchronized 1-day-old adult worms were transferred to the *S. aureus*-seeded plates and incubated at 25°C. Animals were transferred daily to freshly seeded plates, and viability was assessed at the indicated time points. Worms were considered dead if they failed to respond to touch with a platinum wire. For killing assays involving *bar-1* RNAi, worms were grown on empty vector-containing *E. coli* until the L4 stage and subsequently transferred to *bar-1* RNAi plates. The 1-day-old adult worms were then transferred to *S. aureus*-seeded plates and maintained at 25°C. All assays were performed in triplicate with independent biological replicates. Detailed statistical analyses for all survival assays are provided in Table S5.

### *C. elegans* RNA isolation and quantitative reverse transcription-PCR

Wild-type N2 and *usp-14(tm1481)* worms were synchronized by timed egg laying. Approximately 50 gravid adults were transferred to 9-cm NGM plates seeded with either *aex-5* RNAi or control empty vector RNAi bacteria and allowed to lay eggs for 4 hours. The adults were then removed, and the plates were incubated at 20°C for 96 hours. The resulting adult worms were collected, washed with M9 buffer, and preserved in TRIzol reagent (Life Technologies, Carlsbad, CA, USA) for RNA extraction. Total RNA was isolated using the RNeasy Plus Universal Kit (Qiagen, Netherlands). For cDNA synthesis, 1 µg of total RNA was reverse transcribed using random primers and the PrimeScript 1st Strand cDNA Synthesis Kit (TaKaRa), according to the manufacturer’s protocol. Quantitative RT-PCR was performed using TB Green fluorescence (TaKaRa) on a MasterCycler EP Realplex 4 thermal cycler (Eppendorf) in a 96-well plate format. Reactions were prepared in 15 µL volumes and analyzed following the manufacturer’s instructions. Relative transcript levels were calculated using the comparative *CT*(2^−ΔΔ^*CT*) method, normalized to pan-actin (*act-1*, *-3*, *-4*) expression as previously described (Singh & Aballay, 2017). All qRT-PCR reactions were performed in triplicate (technical replicates) and repeated across at least three independent biological replicates. Primer sequences are provided in Table S6.

### RNA sequencing and data analysis

For RNA sequencing, worm synchronization and RNA extraction for N2 and *usp-14(tm1481)* strains grown on *aex-5* RNAi were performed as described above for qRT-PCR. Three independent biological replicates were used for each condition. Library preparation and sequencing were performed at Neuberg Supratech Reference Laboratories, India, with cDNA libraries sequenced on the NovaSeq 6000 platform using 50-bp paired-end reads.

RNA sequencing data were analyzed via the Galaxy web platform (https://usegalaxy.org/) following established protocols (Ghosh & Singh, 2024). Briefly, paired-end reads underwent quality trimming using Trimmomatic before being aligned to the *C. elegans* reference genome (WS220) using STAR. Read counts per gene were obtained using *htseq-count*, and differential gene expression analysis was carried out with DESeq2. Genes showing a ≥2-fold change in expression and a *P*-value < 0.01 were considered differentially expressed. Gene Ontology analysis utilized the DAVID Bioinformatics Database (https://david.ncifcrf.gov/tools.jsp), and Venn diagrams were generated using BioVenn (https://www.biovenn.nl/) (Hulsen *et al*, 2008).

### Plasmid constructs and generation of transgenic *C. elegans*

To generate the *usp-14p::gfp* transcriptional reporter, the 714 bp promoter region upstream of the *usp-14* start codon was amplified from N2 genomic DNA and cloned into the pPD95.77 plasmid upstream of GFP using SalI and KpnI restriction sites. The resulting construct was microinjected into wild-type N2 animals along with pCFJ90 (*myo-2p::mCherry*) as a co-injection marker to generate the *usp-14p::gfp* reporter strain.

For rescuing the *usp-14(jsn19)* mutant phenotype, the *usp-14* promoter and its complete coding sequence, including the stop codon, were amplified from N2 genomic DNA and cloned into pPD95.77 upstream of GFP using SphI and SmaI restriction sites. The construct was microinjected into *usp-14(jsn19);clec-60p::gfp* worms together with pCFJ104 (*myo-3p::mCherry*) as a co-injection marker to generate the strain *usp-14(jsn19);clec-60p::gfp;jsnEx6[usp-14p::usp-14 + myo-3p::mCherry]*.

For neuronal rescue, the 3,392 bp promoter region upstream of the *rgef-1* start codon was first cloned into the pPD95.77 plasmid using BamHI and SmaI restriction sites. The *usp-14* coding sequence was then inserted downstream of the *rgef-1* promoter using the In-Fusion HD Cloning Kit (Takara Bio, Tokyo, Japan). The resulting *rgef-1p::usp-14* construct was microinjected into *usp-14(jsn19);clec-60p::gfp* worms along with pCFJ104 as a co-injection marker to generate *usp-14(jsn19);clec-60p::gfp;jsnEx7[rgef-1p::usp-14 + myo-3p::mCherry]*.

To generate intestine-specific rescue strains, the *usp-14* coding sequence was cloned into the pDB24 plasmid (kindly provided by the laboratory of Alejandro Aballay), which carries the *vha-6* promoter, using BamHI and AgeI restriction sites. The construct was microinjected into *usp-14(jsn19);clec-60p::gfp* worms with pCFJ104 as a co-injection marker to generate *usp-14(jsn19);clec-60p::gfp;jsnEx8[vha-6p::usp-14 + myo-3p::mCherry]*.

For all injections, *usp-14p::gfp*, *usp-14p::usp-14*, *vha-6p::usp-14*, and *rgef-1p::usp-14* plasmids were used at a concentration of 50 ng/µL. The co-injection markers *myo-2p::mCherry* and *myo-3p::mCherry* were used at final concentrations of 5 ng/µL and 25 ng/µL, respectively. All transgenes were maintained as extrachromosomal arrays, and at least two independent transgenic lines were established and analyzed for each construct. Primer sequences are provided in Table S6.

### Fluorescence imaging of *C. elegans*

Fluorescence imaging was performed as described previously (Das *et al*, 2025; Rao *et al*, 2025). Briefly, animals were picked under a non-fluorescence stereomicroscope to avoid potential bias. Worms were anesthetized in M9 buffer containing 50 mM sodium azide and mounted on 2% agarose pads for imaging. Fluorescence visualization was conducted using either a Nikon SMZ-1000 or SMZ18 stereomicroscope. Fluorescence intensity was quantified using ImageJ software. High-resolution images of the *usp-14p::gfp* reporter strain were captured using a ZEISS Imager.Z2 microscope.

### Quantification and statistical analysis

All statistical analyses were performed with Prism 8 (GraphPad). Data are presented as mean ± standard deviation. Statistical significance between two groups was assessed using an unpaired, two-tailed *t*-test. For comparisons involving more than two groups, an ordinary one-way ANOVA followed by Dunnett’s multiple comparisons test was applied. Statistical significance was defined as *p* < 0.05. In all figures, asterisks indicate significance levels as follows: \**p* < 0.05; \*\**p* < 0.01; and \*\*\**p* < 0.001, relative to the corresponding controls. Survival data were analyzed using the Kaplan-Meier method, and differences between survival curves were evaluated using the log-rank test. All experiments were performed independently at least three times unless otherwise stated.

## Acknowledgments

We thank the Caenorhabditis Genetics Center [funded by the NIH Office of Research Infrastructure Programs (P40 OD010440)] for providing the strains used in this study.

## Author Contributions

A.G. and J.S. conceived and designed the experiments. A.G. performed the experiments. A.G. and J.S. analyzed the data and wrote the paper.

## Data availability

The whole-genome sequence data for JSJ19 and JSJ20 have been submitted to the public repository, the Sequence Read Archive, with BioProject ID PRJNA1367285. The RNA sequencing data for N2 and *usp-14(tm1481)* worms grown on the empty vector control and *aex-5* RNAi have been submitted to the public repository, the Sequence Read Archive, with BioProject ID PRJNA1364750. All data generated or analyzed during this study are included in the manuscript and supporting files.

## Supplementary figures

**Figure S1:**
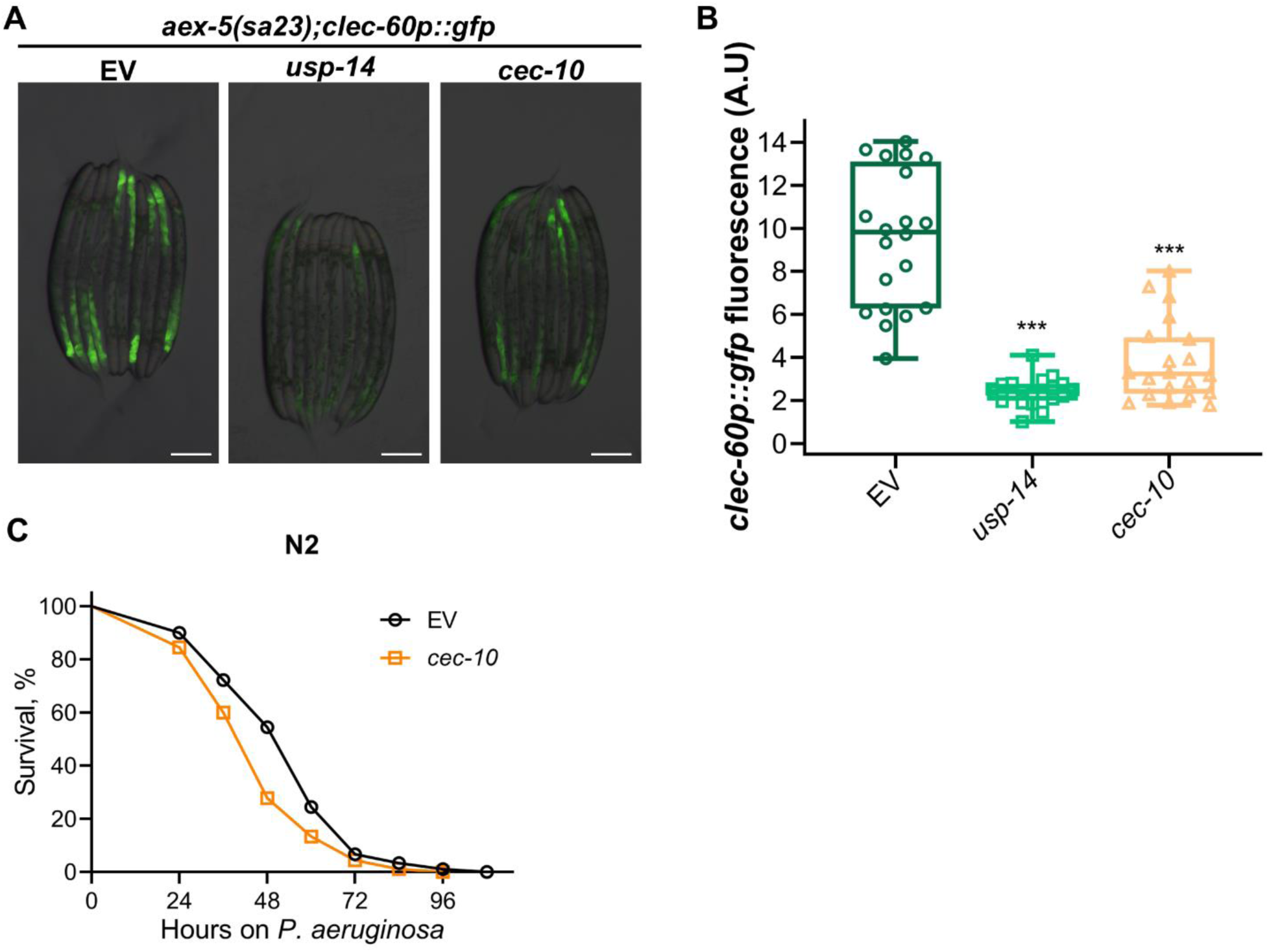
Knockdown of *usp-14* and *cec-10* suppresses intestinal distension-mediated *clec-60* induction. (A) Representative fluorescence images of *aex-5(sa23);clec-60p::gfp* worms grown on the empty vector (EV) control, *usp-14*, and *cec-10* RNAi. Scale bar = 200 μm. (B) Quantification of GFP levels of *aex-5(sa23);clec-60p::gfp* worms grown on the EV control, *usp-14*, and *cec-10* RNAi. \*\*\**p*< 0.001 via ordinary one-way ANOVA followed by Dunnett’s multiple comparisons test (*n* = 20 worms each). (C) Representative survival plots of N2 worms exposed to the EV and *cec-10* RNAi on *P. aeruginosa* PA14 at 25°C. \*\*\**p*< 0.001 (*n* = 60 on EV and 90 on *cec-10* RNAi).

**Figure S2:**
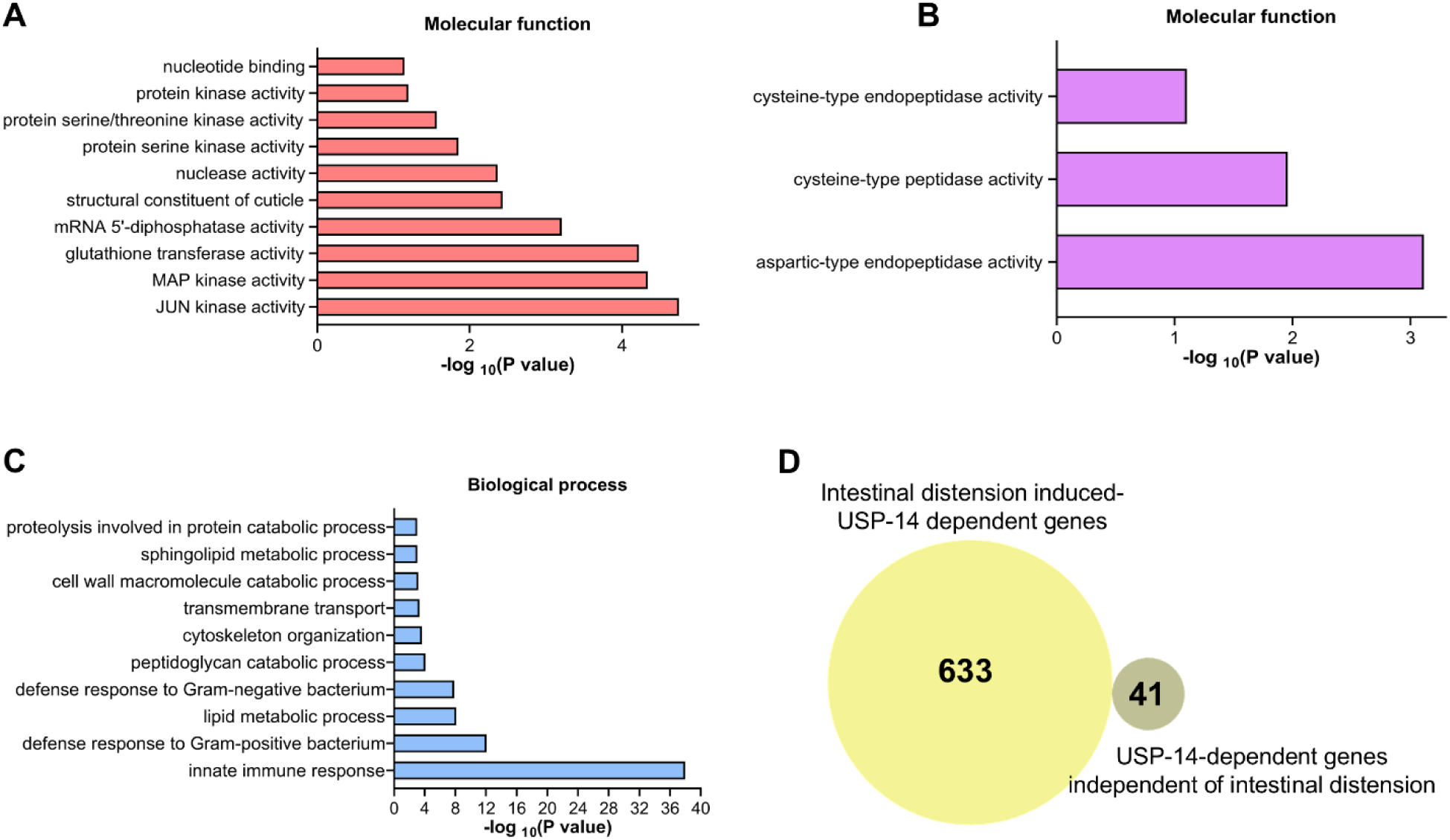
Intestinal distension induces USP-14-dependent and -independent innate immune response genes. (A)-(B) Gene Ontology (GO) enrichment analysis for molecular functions for upregulated (A) and downregulated (B) genes in *usp-14(tm1481)* worms compared to wild-type N2 worms grown on the empty vector (EV) control RNAi. (C) GO enrichment analysis for biological process for upregulated genes in N2 worms grown on *aex-5* RNAi compared to N2 worms grown on the EV control RNAi. (D) Venn diagram showing the overlap between intestinal distension-induced USP-14-dependent genes and genes downregulated in *usp-14(tm1481)* mutants independent of intestinal distension.

**Figure S3:**
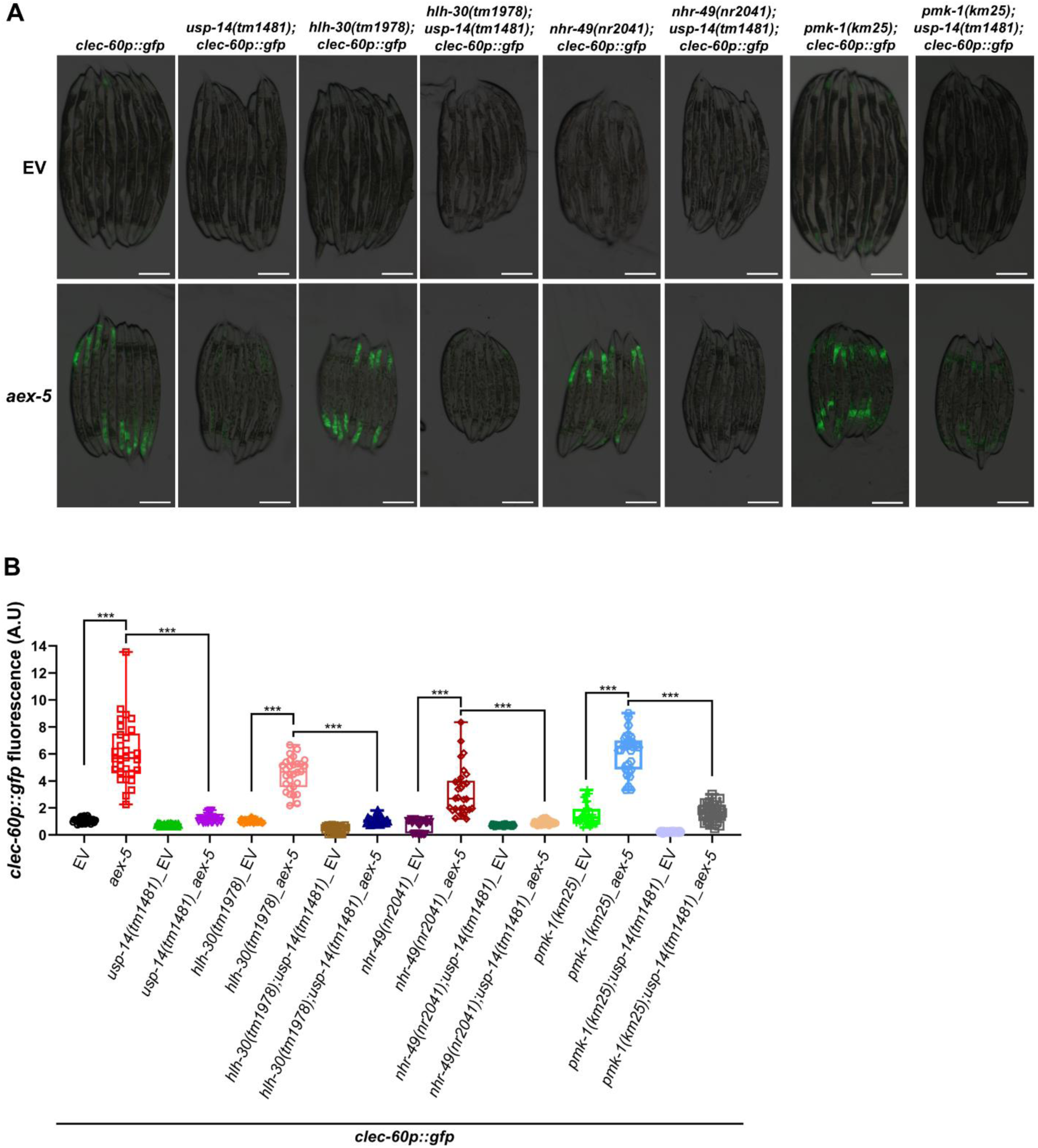
USP-14 mediates intestinal distension-induced immunity independent of known canonical innate immune pathways. (A) Representative fluorescence images of *clec-60p::gfp*, *usp-14(tm1481);clec-60p::gfp, hlh-30(tm1978);clec-60p::gfp, hlh-30(tm1978);usp-14(tm1481);clec-60p::gfp*, *nhr-49(nr2041);clec-60p::gfp*, *nhr-49(nr2041);usp-14(tm1481);clec-60p::gfp*, *pmk-1(km25);clec-60p::gfp*, and *pmk-1(km25);usp-14(tm1481);clec-60p::gfp* worms exposed to empty vector (EV) control and *aex-5* RNAi. Scale bar = 200 μm (B) Quantification of GFP levels of *clec-60p::gfp*, *usp-14(tm1481);clec-60p::gfp, hlh-30(tm1978);clec-60p::gfp, hlh-30(tm1978);usp-14(tm1481);clec-60p::gfp*, *nhr-49(nr2041);clec-60p::gfp*, *nhr-49(nr2041);usp-14(tm1481);clec-60p::gfp*, *pmk-1(km25);clec-60p::gfp*, and *pmk-1(km25);usp-14(tm1481);clec-60p::gfp* worms exposed to EV control and *aex-5* RNAi. \*\*\**p*< 0.001 via unpaired t-test (*n* = 30 worms each).

**Figure S4:**
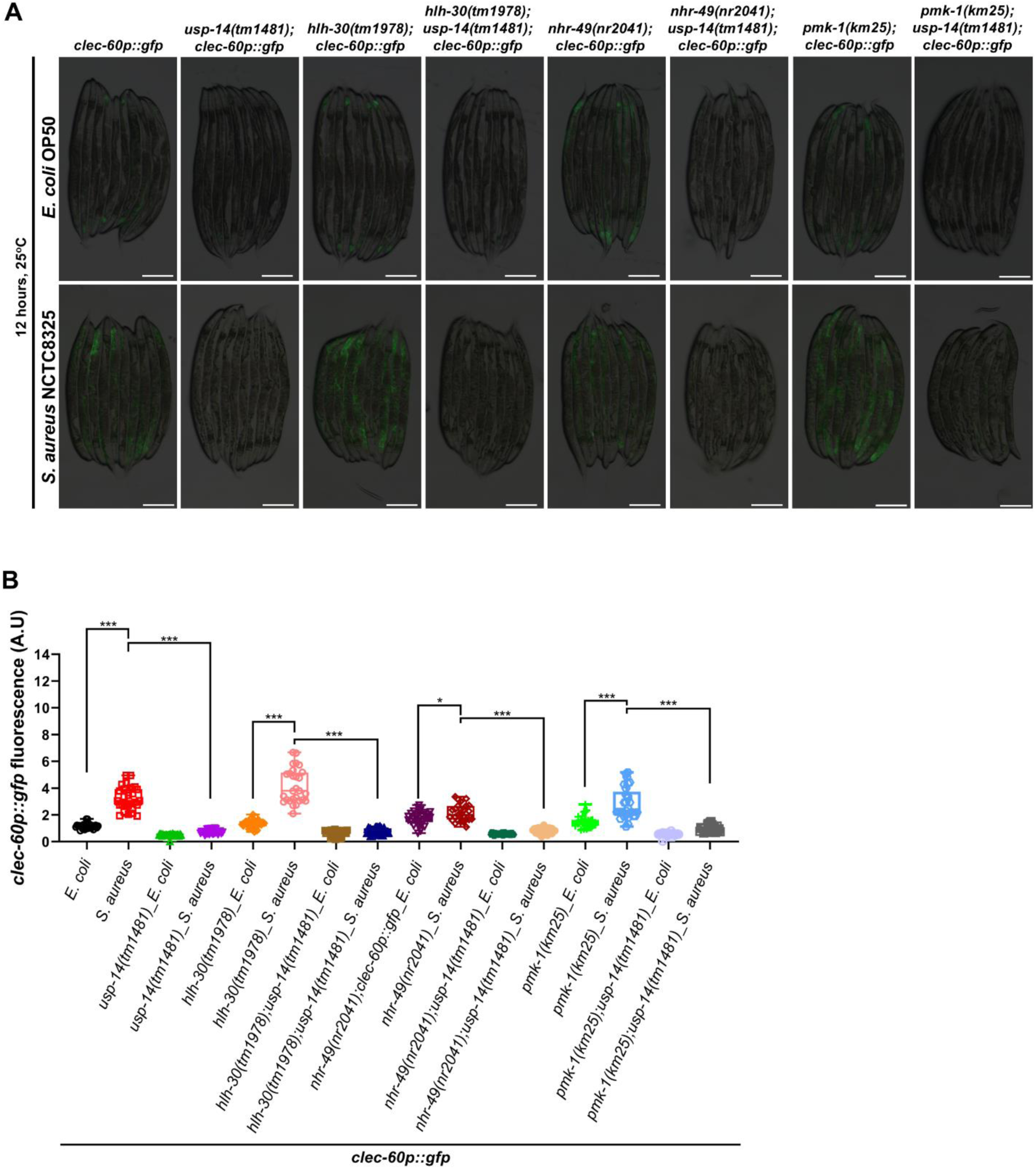
USP-14 mediates *S. aureus*-induced immunity independent of known canonical innate immune pathways. (A) Representative fluorescence images of *clec-60p::gfp*, *usp-14(tm1481);clec-60p::gfp, hlh-30(tm1978);clec-60p::gfp, hlh-30(tm1978);usp-14(tm1481);clec-60p::gfp*, *nhr-49(nr2041);clec-60p::gfp*, *nhr-49(nr2041);usp-14(tm1481);clec-60p::gfp*, *pmk-1(km25);clec-60p::gfp*, and *pmk-1(km25);usp-14(tm1481);clec-60p::gfp* worms after 12 hours of infection with *S. aureus* NCTC8325, compared with *E. coli* OP50 controls at 25 °C. Scale bar = 200 μm (B) Quantification of GFP levels of *clec-60p::gfp*, *usp-14(tm1481);clec-60p::gfp, hlh-30(tm1978);clec-60p::gfp, hlh-30(tm1978);usp-14(tm1481);clec-60p::gfp*, *nhr-49(nr2041);clec-60p::gfp*, *nhr-49(nr2041);usp-14(tm1481);clec-60p::gfp*, *pmk-1(km25);clec-60p::gfp*, and *pmk-1(km25);usp-14(tm1481);clec-60p::gfp* worms after 12 hours of infection with *S. aureus* NCTC8325, compared with *E. coli* OP50 controls at 25 °C. \*\*\**p*< 0.001 and \**p*< 0.05 via unpaired t-test (*n* = 27-32 worms each).

**Figure S5:**
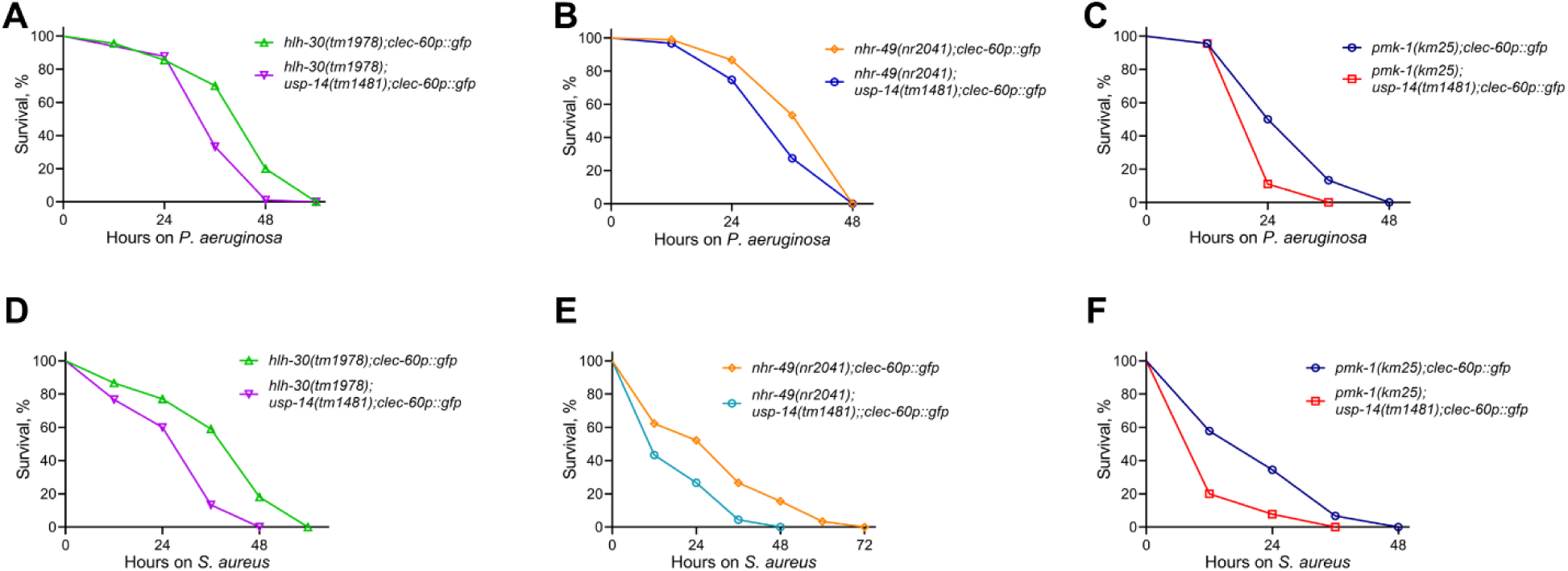
USP-14 modulates immunity independent of known canonical innate immune pathways. (A) Representative survival plots of *hlh-30(tm1978);clec-60p::gfp* and *hlh-30(tm1978);usp-14(jsn19);clec-60p::gfp* worms on *P. aeruginosa* PA14 at 25°C. \*\*\**p*< 0.001 (*n* = 90 per condition). (B) Representative survival plots of *nhr-49(nr2041);clec-60p::gfp* and *nhr-49(nr2041);usp-14(tm1481);clec-60p::gfp* worms on *P. aeruginosa* PA14 at 25°C. \*\*\**p*< 0.001 (*n* = 90 for *nhr-49(nr2041);clec-60p::gfp* and 91 for *nhr-49(nr2041);usp-14(tm1481);clec-60p::gfp*). (C) Representative survival plots of control *pmk-1(km25);clec-60p::gfp* and *pmk-1(km25);usp-14(tm1481);clec-60p::gfp* worms on *P. aeruginosa* PA14 at 25°C. \*\*\**p*< 0.001 (*n* = 90 per condition). (D) Representative survival plots of control *hlh-30(tm1978);clec-60p::gfp* and *hlh-30(tm1978);usp-14(jsn19);clec-60p::gfp* worms on *S. aureus* NCTC8325 at 25°C. \*\*\**p*< 0.001 (*n* = 90 per condition). (E) Representative survival plots of control *nhr-49(nr2041);clec-60p::gfp* and *nhr-49(nr2041);usp-14(tm1481);clec-60p::gfp* worms on *S. aureus* NCTC8325 at 25°C. \*\*\**p*< 0.001 (*n* = 90 per condition). (F) Representative survival plots of control *pmk-1(km25);clec-60p::gfp* and *pmk-1(km25);usp-14(tm1481);clec-60p::gfp* worms on *S. aureus* NCTC8325 at 25°C. \*\*\**p*< 0.001 (*n* = 90 per condition).

## Supplementary tables

**Table S1:** Upregulated and downregulated genes in *aex-5* RNAi versus empty vector control RNAi in N2 worms. Genes exhibiting at least two-fold change and *P*-value <0.01 were considered differentially expressed. A separate Excel file is provided.

**Table S2:** Upregulated and downregulated genes in *aex-5* RNAi versus empty vector control RNAi in *usp-14(tm1481)* worms. Genes exhibiting at least two-fold change and *P*-value <0.01 were considered differentially expressed. A separate Excel file is provided.

**Table S3:** Upregulated and downregulated genes in *usp-14(tm1481)* worms versus N2 worms grown on empty vector control RNAi. Genes exhibiting at least two-fold change and *P*-value <0.01 were considered differentially expressed. A separate Excel file is provided.

**Table S4:** Comparison of genes upregulated upon *aex-5* RNAi in N2 and *usp-14(tm1481)* worms. A separate Excel file is provided.

**Table S5:** Statistical analysis of survival curves from three independent experiments for each condition. A separate Excel file is provided.

**Table S6:**
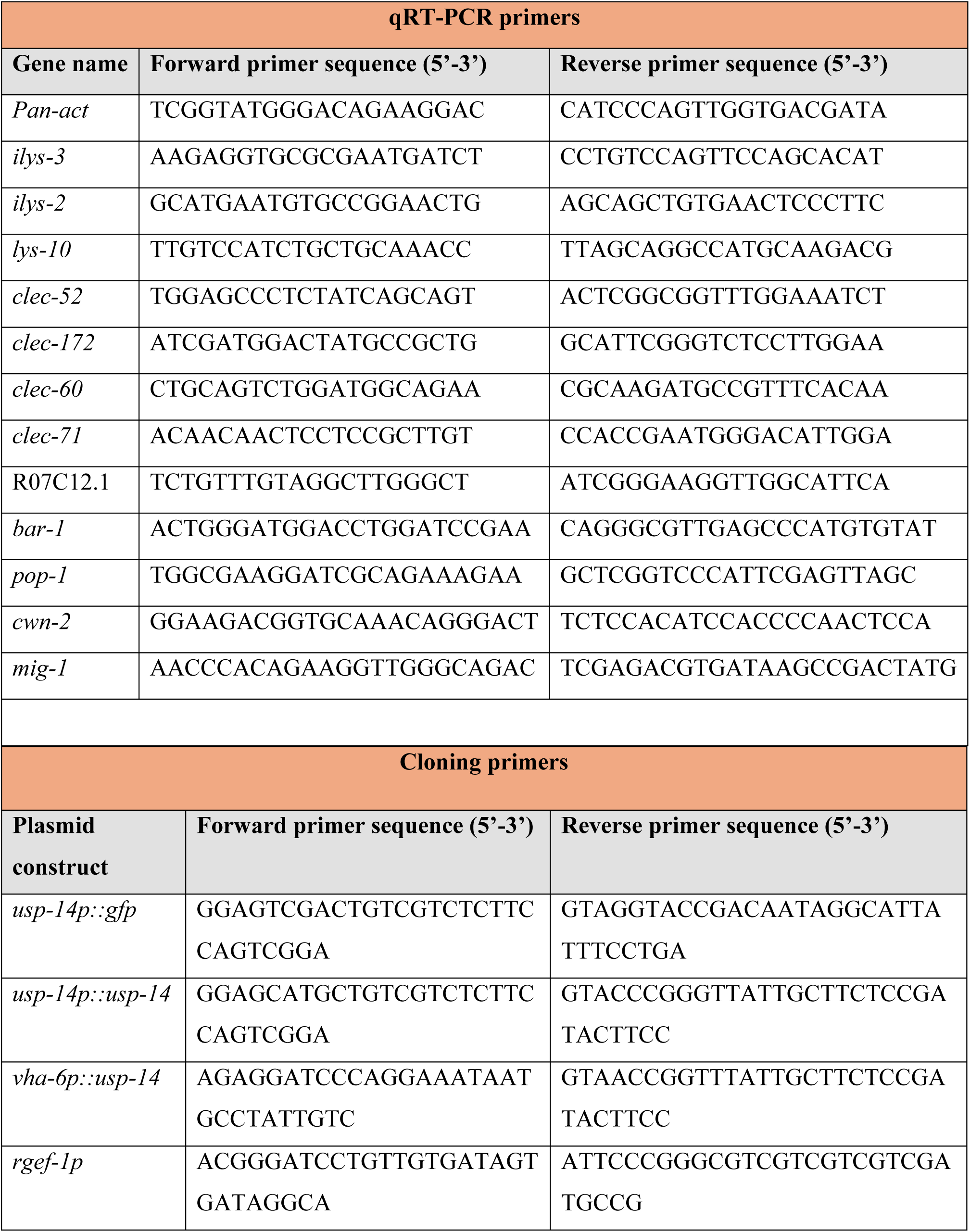
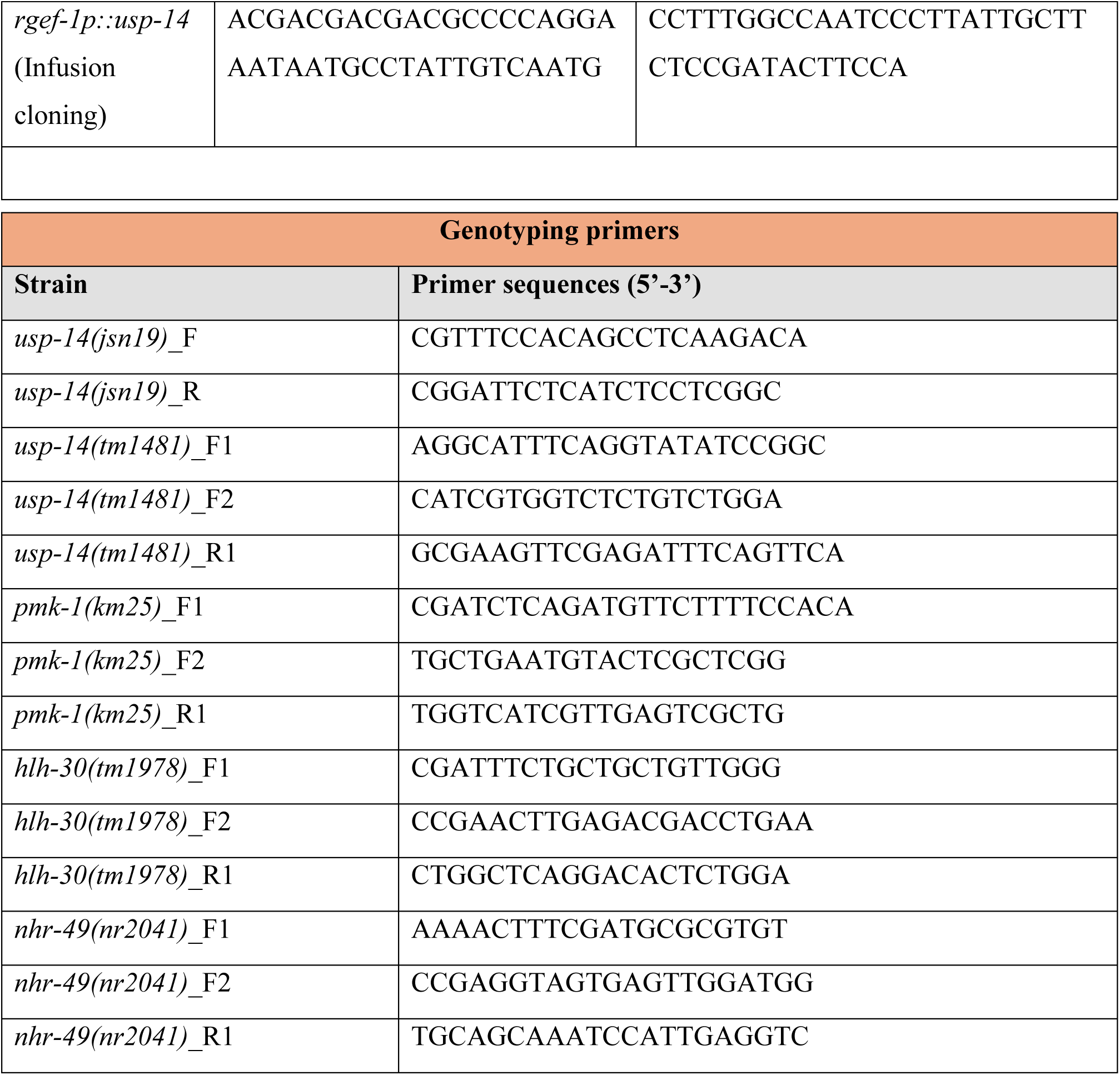
Primers used in the study.

